# Functional specialization of *TAF4b* governs early prophase I entry and recombination frequency in *Brassica rapa*

**DOI:** 10.64898/2026.09.29.754957

**Authors:** Joanna Majka, Zhenling Lv, Thu Dieu Nguyen, Rebecca Doll, Jan Schonenbach, Annaliese S. Mason

## Abstract

While the core transcription factor *TAF4b* regulates meiotic transcription and recombination in the diploid model *Arabidopsis thaliana*, its functional specialization and evolutionary dynamics remain poorly understood in paleopolyploid genomes. Here, we investigated how a unique, single-copy *TAF4b* gene on chromosome A7 modulates early meiotic progression and reproductive success in the paleopolyploid crop *Brassica rapa*. By characterizing multiple independent *taf4b* TILLING mutant lines through an integrated pipeline of whole-genome background sequencing, stage-specific anther transcriptomics (RNA-seq), yeast two-hybrid assays, and quantitative cytogenetics, we show that *TAF4b* has evolved an exclusive, non-redundant, and meiosis-enriched role despite the presence of duplicated somatic *TAF4 loci*. Disruption of *TAF4b* universally reduced class I crossover frequency, critically depleted HEI10 crossover-designation foci, and triggered univalent formation leading to severe fertility loss. Our stage-specific RNA-seq profiling and structural modeling demonstrate that this phenotype is driven by a precise transcriptomic dysregulation strictly restricted to the narrow interphase/leptotene transition. Crucially, this early transcriptomic imbalance disrupts the expression stoichiometry of core axis, synapsis, and recombination regulators, including ASY1, REC8, SWI1, and BRCA2B, leading to prolonged ASY1 retention and defective physical chromosome behavior. In conclusion, we demonstrate that *TAF4b* acts as a stage-specific upstream gatekeeper of the early prophase I expression program in *B. rapa*. These findings establish a direct link between core transcriptional gating, downstream chromosome dynamics, and meiotic stability following evolutionary genome duplication.

## Introduction

Meiotic recombination is essential for both faithful chromosome segregation and the generation of genetic diversity in sexually reproducing organisms. During prophase I of meiosis, programmed double-strand breaks (DSBs) are repaired through homologous recombination, resulting in either crossovers (COs) or non-crossovers (gene conversion). COs, which manifest cytologically as chiasmata, physically link homologous chromosomes and ensure their correct segregation in the first meiotic division (Barton 2009; Zickler & Kleckner 2015; Lambing et al. 2017). In plants, CO number is typically tightly constrained by the obligatory crossover and by strong crossover interference, resulting in a limited number of COs per chromosome pair despite the formation of many DSBs (Mercier et al. 2015; Lloyd & Bomblies 2016). This inherent limitation on recombination restricts the ability to reshuffle genetic variation and remains a major barrier for crop improvement. Consequently, identifying the factors that either limit or stimulate CO formation is critical for understanding and modulating the meiotic recombination landscape.

Considerable progress has been made in identifying the molecular pathways controlling CO formation. Two main classes of COs are recognized in plants: interference-sensitive class I COs, which depend on a conserved ZMM protein pathway, and interference-insensitive class II COs, which are resolved primarily by structure-specific endonucleases (Higgins et al. 2004; Mercier et al. 2015). In addition to canonical meiotic factors, an increasing number of chromatin-associated and transcription-related proteins have been implicated in modulating recombination frequency and distribution (Yelina et al. 2015; Lambing et al. 2020). These findings indicate that CO control is intimately linked to broader regulatory process, such as chromatin organization and transcriptional control.

One genetic factor implicated in CO formation is *TAF4b* (TATA-box binding protein-associated factor 4B), a germline-enriched subunit of the TFIID transcription initiation complex. In animals, *TAF4b* is required for proper germ cell development and fertility (Freiman et al. 2001; Falender et al. 2005; Grive et al. 2016). In plants, *Arabidopsis thaliana TAF4b* has been shown to play important roles in reproductive development and meiotic progression, but its loss does not significantly reduce overall fertility or result in sterility (Lawrence et al. 2019). Genome-wide recombination analyses in *A. thaliana* suggested that mutation of *TAF4b* leads to a reduction in CO frequency (decreases ranging from 9.0% to 27.1%), mostly in sub-telomeric regions (Lawrence et al. 2019).

Whether the role of *TAF4b* in crossover regulation is conserved across species, or how evolutionary dynamics shape its functional landscape in polyploids, remains unknown. This question is particularly relevant in the genus *Brassica*, which comprises several globally important crops and exhibits complex genome evolution following whole-genome duplication events (Cheng et al. 2016). Although *Brassica rapa* underwent a paleopolyploid history (mesohexaploid), it has retained only a single copy of *TAF4b* on chromosome A7, alongside duplicated copies of its related counterpart, *TAF4* (Cheng et al. 2013). As a crop relative of *A. thaliana* in the Brassicaceae family, *B. rapa* serves as a powerful model system for the comparative analysis of meiotic recombination (The *Brassica rapa* Genome Sequencing Project Consortium 2011; Lloyd et al. 2018). Despite extensive synteny between the two genomes, recombination landscapes in *B. rapa* differ markedly from those of *A. thaliana,* reflecting divergent chromatin organization, structural variation, and centromere architecture (Lloyd et al. 2018). These differences position the *B. rapa TAF4b* locus as a compelling system for future studies addressing how core transcription factors maintain non-redundant roles in an agronomically relevant context.

The primary objective of this study was to elucidate how the unique *TAF4b* gene modulates the meiotic landscape within the complex genome of *B. rapa*. We hypothesized that post-polyploidization genome remodeling led to distinct functional specialization between *TAF4* and *TAF4b*, with the single-copy *TAF4b* gene on chromosome A7 acting as a non-redundant regulator of the early meiotic program. We leveraged a *B. rapa* TILLING population as a reverse-genetics resource to identify a comprehensive allelic series targeting distinct domains of this core factor. Here, we systematically characterized multiple independent *taf4b* mutant variants, focusing on a highly conserved functional cluster, and assessed their impact on early meiotic progression, crossover frequency, and reproductive success using integrated quantitative cytogenetics, biochemical interaction assays, and high-throughput sequencing. In this work, we demonstrate that structural disruption of this unique *TAF4b* locus uncovers a novel, non-redundant role in regulating early meiotic entry and crossover assurance that differs distinctively from the phenotype reported in *A. thaliana*. By characterizing these transcriptional, biochemical, and cytogenetic alterations, our study provides clear evidence of how the evolutionary and functional divergence of core transcription factors governs key reproductive processes in an agronomically important crop species.

## Materials and Methods

### Plant Material

*Brassica rapa* mutant lines were obtained from RevGenUK (https://www.jic.ac.uk/research-impact/technology-research-platforms/reverse-genetics/): *taf4b*_1 (JI40382-A), *taf4b*_2 (JI30215-B), *taf4b*_3 (JI30511-B), *taf4b*_4 (JI31320-B), *taf4b*_5 (JI30848-B), and *B. rapa* cultivar R-o-18 (used as the wild-type (WT)). For each *taf4b* mutant line, five seeds were initially sown. Due to incomplete germination, early plant death, or failure to flower, an additional six seeds were sown for *taf4b*_1. All plants were grown in a controlled climate chamber under 16 h light/8 h dark at 21 °C / 18 °C day/night temperatures.

### DNA extraction and genotyping

Genomic DNA was extracted using the CTAB method (Doyle & Doyle 1987). PCR-based genotyping was performed using Primer3-designed primers and *Taq* DNA polymerase in a 25 µL reaction volume. The PCR conditions consisted of 32 cycles with an optimized annealing temperature of 57-59 °C. Finally, the PCR products were submitted to Eurofins Genomics for Sanger sequencing. Information regarding mutation types, sequencing results, and primer sequences is shown in Table 1. DNA samples sent for whole genome sequencing were extracted using the Plant DNA Kit (VWR) according to the provided manual.

**Table 1.**
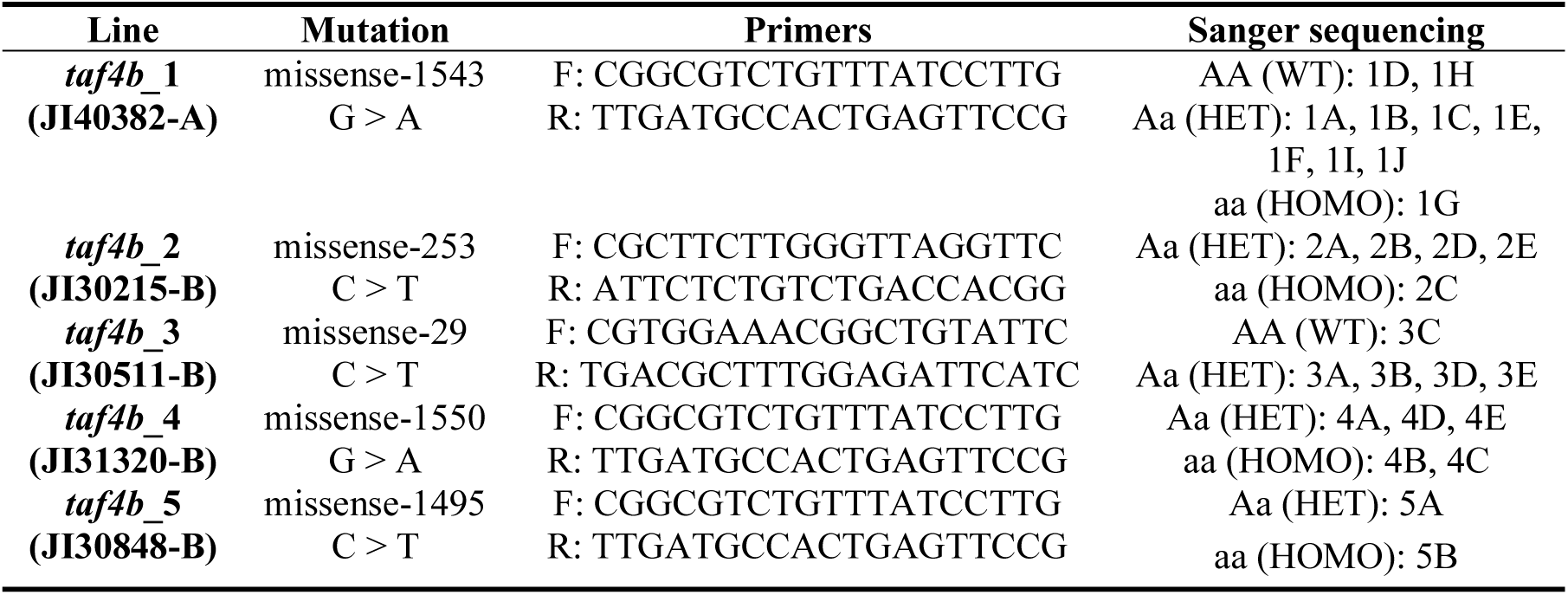
Characteristics of TILLING lines, primer sequences, and Sanger sequencing-based genotyping results for the analyzed *B. rapa* plants.

### Pollen viability

Pollen viability was assessed using 1% acetocarmine staining (see in Addo Nyarko et al. 2024). For each plant, three open flowers from different branches were collected. After staining, pollen grains were imaged using an Olympus Axiolab 5 microscope. Regularly shaped, pink-stained pollen grains were scored as viable, whereas small, irregularly shaped, yellow/unstained or shrunken grains were scored as non-viable. At least 900 pollen grains were counted from each plant.

### Chromosome preparations for chiasma counting

Inflorescences were collected from approximately nine-week-old *B. rapa* plants and fixed in Carnoy’s solution (3:1, 99.8% ethanol: glacial acetic acid) for 24 hours at room temperature. After fixation, flower buds were transferred to 70% ethanol and stored at −20°C. Prior to slide preparation, flower buds were staged by examining one anther per bud. Anthers were digested in an enzyme solution containing 5% (w/v) cellulase Onozuka R10 and 1% (w/v) pectolyase Y23 for 80 minutes at 37°C. Chromosome slides were prepared using the dropping technique and counterstained with DAPI in Vectashield (Vector Laboratories). For each plant, we took and scored pollen mother cells that came from at least two buds from two different branches.

### Immunolocalization

Anthers at pachytene, diakinesis, metaphase I were digested in an enzyme mixture consisting of 0.3% (w/v) cellulase Onozuka R10, 0.3% (w/v) pectolyase *Aspergillus japonicus*, and 0.3% (w/v) from *Helix pomatia* for 3 hours and 30 minutes at 37°C. After digestion, anthers were dissected, chopped and squeezed in 1% acetocarmine, and slides were frozen in liquid nitrogen. Slides containing appropriate meiotic stages were selected based on DAPI counterstaining. Selected slides were treated with 100 μg/ml RNase A in 1 × PBS (Sigma Aldrich) for 60 min. Slides were washed in 1 × PBS buffer and incubated for 48 - 72 hours with a primary rabbit anti-HEI10 antibody (1:100 dilution), anti-guinea pig ASY1 (1:100), and anti-rabbit ZYP1 (1:150). Subsequently, slides were rinsed three times in 1 × PBS with 0.1% Triton X100 (PBST) (Sigma Aldrich), incubated with an appropriate secondary antibody conjugated with a fluorescent dye (Alexa 488, Alexa594 and Alexa647) for 1 hour and 30 minutes at 37°C and washed again three times in 1 × PBST.

All slides were examined using an Axio Observer 7 inverted microscope (Zeiss, Germany) equipped with an Axiocam 305 camera (Zeiss, Germany). Images were acquired using Zeiss ZEN v3.4 software and then processed using Adobe Photoshop CS5 software. To investigate synapsis, images were acquired using a Zeiss LSM 980 confocal microscope equipped with an Airyscan 2 detector (Zeiss, Germany) operated in super-resolution (SR) mode. Airyscan image reconstruction and analysis were performed in ZEN software (v.3.13; Zeiss) using default SR processing settings.

### Transcriptome sequencing and differential expression analysis

Total RNA was extracted from young leaves and fresh anthers at specific meiotic stages (premeiosis/interphase, pachytene, and telophase II) using the Monarch Total RNA Miniprep Kit (New England Biolabs) with on-column DNase I treatment. RNA quality was verified on a Bioanalyzer 2100 (Agilent Technologies, Santa Clara, CA, USA), and high-integrity samples were selected for paired-end library preparation. Sequencing was performed on an Illumina NovaSeq 6000 platform (Illumina Inc., San Diego, CA, USA), yielding 2807 million 150-bp reads. Raw reads were quality-trimmed using Trimmomatic with a sliding-window approach (Q20), discarding adapter sequences and reads shorter than 50 bp. Clean reads were aligned to the reference genome via HISAT2 (Supplementary Table S1). Gene counts generated by StringTie and featureCounts were analyzed using DESeq2, defining differentially expressed genes (DEGs) by an adjusted P-value < 0.05 and |log₂FC| ≥ 1.5.

### Variant discovery, annotation, and candidate filtering

Genomic DNA was extracted from young leaves using a DNeasy Plant Mini Kit (Qiagen) and sequenced using the Illumina NovaSeq 6000 platform to generate 1139 million 150-bp paired-end reads. High-quality reads were mapped to the *Brassica rapa* reference genome (R-o-18 v2.3) using BWA-MEM (Li et al. 2013) (Supplementary Table S2). After removing duplicates, variants were called and filtered using GATK (VariantFiltration) and bcftools (Danecek et al. 2021). Standard hard-filtering (MQ < 40, QUAL < 30, QD < 2.0, FS > 60, SOR > 3) and subsequent refinement (QUAL/MQ ≥ 40, DP ≥ 10, QD ≥ 2.0, FS ≤ 30, SOR ≤ 3) yielded 148 443 high-confidence SNPs after excluding sites with low polymorphism (< 3 individuals carrying the alternative allele). Variants were annotated using SnpEff (Cingolani et al. 2012), identifying 925 high-impact and 6 265 moderate-impact mutations. Putative protein phosphorylation sites generated or disrupted by the identified amino acid substitutions were predicted in silico using the NetPhos 3.1 server (Blom et al. 2004) with default thresholds. To eliminate background mutations from the TILLING lines, variants present in the wild-type reference were removed, retaining only loci where mutant genotypes (Aa/aa) differed from the homozygous wild-type (AA). Candidate SNPs were subsequently validated via PCR and Sanger sequencing.

### Molecular cloning work for yeast two-hybrid (Y2H) construction

Total RNA was extracted from the young leaf tissues of *B. rapa* R-o-18 (WT) and the *taf4b*_1_aa mutant line (JI40382-A) using the Monarch Total RNA Miniprep Kit (New England Biolabs), including an on-column DNase I treatment. First-strand cDNA was synthesized from 2 µg of total RNA using oligo(dT)_18_ primers and the RevertAid First Strand cDNA synthesis kit (Thermo Fisher Scientific) according to the manufacturer’s instructions. Full-length open reading frames (ORFs) of *TAF4b*-WT (A07p016600.1_BraROA.1), mutated *TAF4b* (TAF4b-M) and *TAF12* (*A05p049220.1_BraROA.1*) were amplified from cDNA using specific primers (Supplementary Table S3) and the Q5® Hot Start High-Fidelity DNA Polymerase (New England Biolabs).

The resulting PCR products were adapted with Gateway attB flanking sequences and cloned into the Gateway™ pDONR™/Zeo entry vector (Invitrogen) via BP clonase reactions. Plasmids were propagated in *Escherichia coli* strain 5-alpha and verified by whole-plasmid Nanopore sequencing (Eurofins Genomics). To generate bait and prey constructs for the yeast two-hybrid (Y2H) assay, the verified ORFs were transferred via Gateway LR recombination reactions into either pDEST™32 (to generate GAL4 DNA-binding domain [DBD] fusions with a LEU2 marker) or pDEST™22 (to generate GAL4 activation domain [AD] fusions with a TRP1 marker). All destination constructs were confirmed by Nanopore sequencing.

### Yeast transformation and autoactivation/toxicity assays

Yeast transformation was performed using the ProQuest™ Two-Hybrid System (Invitrogen) in the *Saccharomyces cerevisiae* reporter strain *MaV203*, following a standard lithium acetate/single-stranded carrier DNA/polyethylene glycol (LiAc/SS-DNA/PEG) protocol. Transformants were selected on synthetic complete (SC) medium lacking leucine (SC-Leu) for bait-only constructs, or lacking leucine and tryptophan (SC-Leu-Trp) for bait-prey co-transformations, and incubated at 30°C for 3 days. Plasmid combinations are detailed in Supplementary Table S4. To evaluate bait toxicity, growth rates of yeast expressing pDEST32::TAF12, pDEST32::TAF4b-WT, or pDEST32::TAF4b-M were compared to an empty pDEST32 control. Autoactivation potential was assessed by co-transforming each bait plasmid with an empty pDEST22 prey vector (Supplementary Table S4). The pEXP™32/Krev1 plasmid co-transformed with pEXP™22/RalGDS-wt, -m1, or -m2 served as positive and negative controls, alongside an empty pDEST32/pDEST22 vector control. Overnight liquid cultures were normalized, spotted (5 µL) onto solid SC-Leu-Trp-Ura selection medium, and grown at 30°C for 2 days.

### Yeast two-hybrid screening

Y2H screening was executed according to the ProQuest™ system guidelines with minor modifications. All media were supplemented with 50 µg/mL kanamycin to prevent bacterial contamination. For the β-galactosidase assay, X-Gal was added directly to YPAD medium to a final concentration of 80 µg/mL. Normalized yeast cultures were spotted (5 µL) onto SC-Leu-Trp control plates and interaction selection media, which included SC-Leu-Trp-Ura, SC-Leu-Trp-His supplemented with 3-amino-1,2,4-triazole (3-AT; 0, 10, 25, 50, and 100 mM), SC-Leu-Trp supplemented with 5-fluoroorotic acid (5-FOA; 0.1% and 0.2%), and X-Gal selection plates.

## Results

### Identification of the unique *B. rapa TAF4b* gene and selection of mutant lines

BLAST-based homology searches identified a single *TAF4b* gene in *B. rapa* located on chromosome A7 (BraA07g012590.4.1C), alongside two copies of its related counterpart, *TAF4*, located on chromosomes A2 and A6 (BraA02g031760.4.1C and BraA06g046780.4.1C, respectively). To determine whether these distinct loci exhibit developmental subfunctionalization, we compared normalized RNA-seq counts retrieved from somatic leaf tissues and staged meiotic anthers. While the *TAF4*-A2 and *TAF4*-A6 loci displayed broadly baseline, static expression levels between leaves and meiotic samples, the *TAF4b*-A7 gene showed significant upregulation specifically during the interphase/leptotene transition relative to somatic tissues (Fig. 1 and Supplementary Table S5).

**Figure 1.**
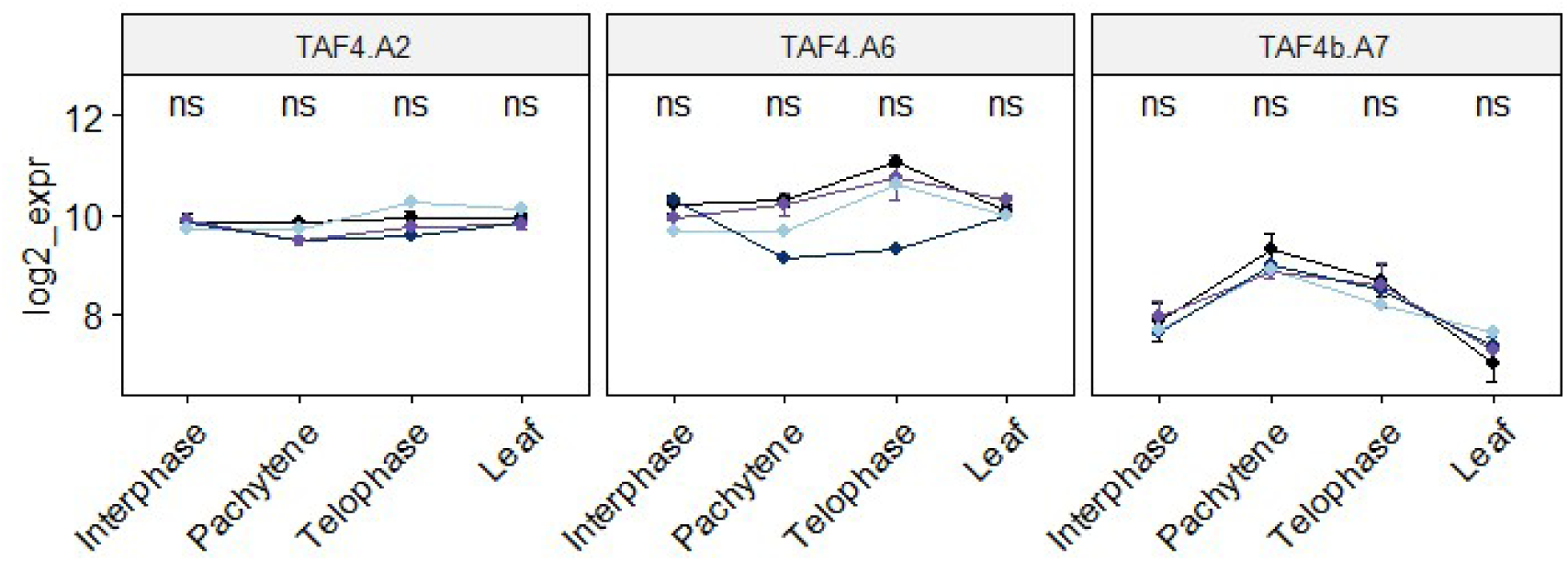
Meiosis-enriched expression of the single-copy *TAF4b* gene on chromosome A7 in *Brassica rapa*. Relative expression levels (log_2_-transformed) of *TAF4b* located on chromosome A7, and its related *TAF4* counterparts (located on chromosomes A2 and A6) across staged anthers (interphase, pachytene, telophase) and vegetative leaf tissue. Line colors represent distinct genetic backgrounds: wild-type R-o-18 (black), *taf4b*_aa (light blue), *taf4b*_Aa (purple), and *taf4b*_AA (navy). ‘ns’ indicates no significant difference in expression levels between the genotypes at the corresponding developmental stage (Tukey’s HSD post-hoc test (*P* > 0.05).

This distinct, meiosis-enriched expression profile identified the chromosome A7 copy as the primary candidate driving early reproductive transcription. Five independent mutant lines (*taf4b*_1 to *taf4b*_5) for targeting this specific gene were acquired from the John Innes Centre *B. rapa* TILLING genomic resource (Table 1; Fig. 2a). Sanger sequencing confirmed that all five lines carry missense substitutions at distinct coding positions within the A7 *TAF4b* sequence (Table 1). During reproductive development, several lines exhibited noticeable abnormalities, including reduced flower bud production and occasional inflorescence abortion, which resulted in shorter siliques or aborted siliques and defective pollen grains (Fig. 2b-e).

**Figure 2.**
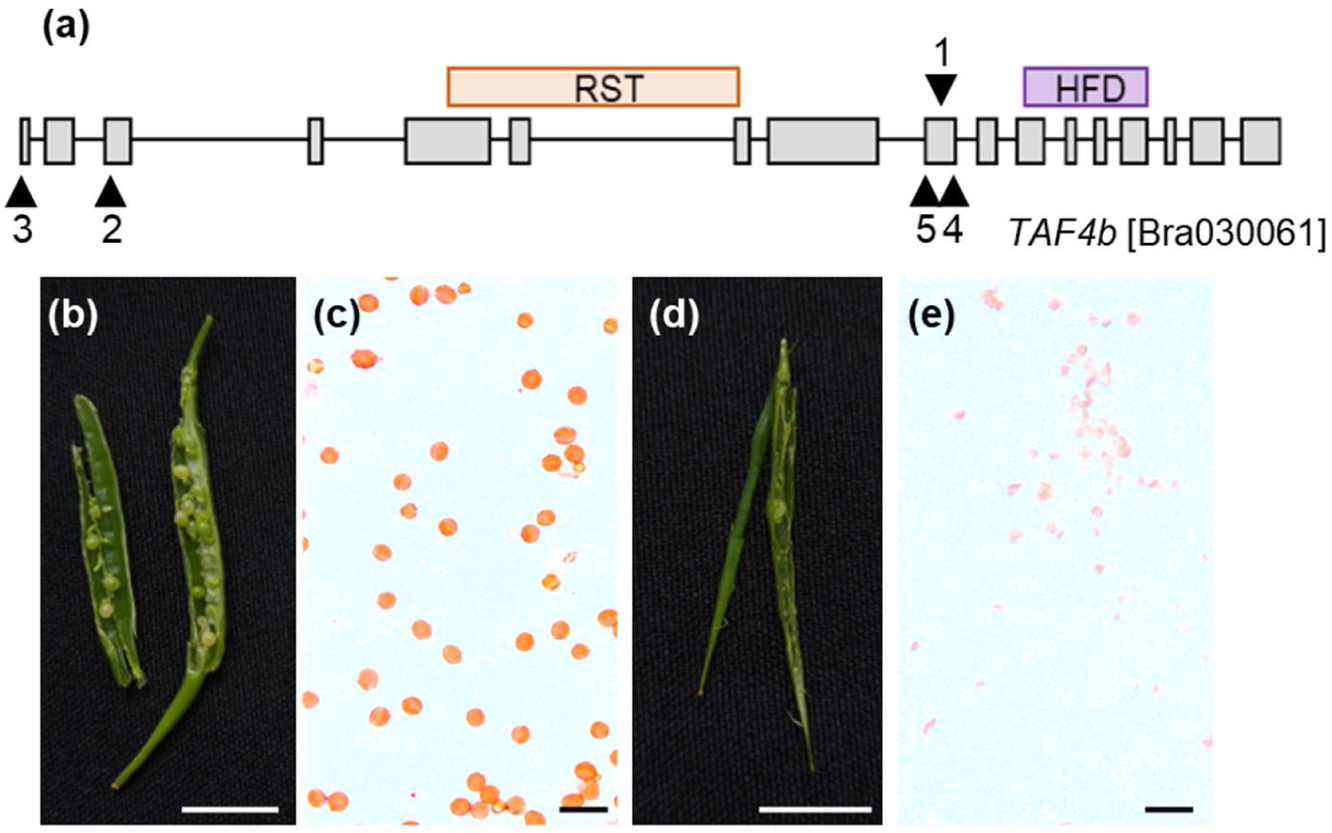
Structural domain architecture of the *TAF4b* locus (Bra030061) and reproductive phenotypes of *Brassica rapa* wild-type (WT, R-o-18) and the *taf4b*_1 mutant. **(a)** Schematic diagram illustrating the exon-intron structure of the *TAF4b* gene and predicted protein domains. Gray rectangles indicate exons, lines indicate introns, and colored horizontal bars denote the boundaries of the RST (aa 224-295) and HFD (aa 584-608) domains. Arrowheads with numbers 1-5 mark the physical position of mutations in the respective TILLING lines (detailed in Table 1). Silique morphology of a WT plant **(b)** and the *taf4b*_1 mutant **(d)**. Pollen viability assessment in WT plant **(c)** and the *taf4b*_1 mutant **(e)**. Scale bars: 1 cm (b, d); 100 µm (c, e).

### Disruption of the *TAF4b* locus universally reduces meiotic crossover frequency

Our initial cytological survey of male meiotic progression revealed that chromosome behavior was largely conventional across most mutant lines. However, distinct meiotic defects were detected in the *taf4b* mutant, characterized by the presence of univalents at diakinesis (Supplementary Fig. S1) that persisted into metaphase I, causing unequal chromosome segregation at telophase I and telophase II (Fig. 3). Despite the globally regular meiotic progression in the remaining lines, a detailed evaluation of chromosome configurations at diakinesis uncovered general modifications in bivalent morphology across the entire allelic series, manifested by a higher ratio of rod bivalents relative to ring bivalents. To quantify this potential shift in crossover frequency, we scored physical chiasmata numbers at the diakinesis/metaphase I stage. Wild-type *B. rapa* cells displayed a mean of 17.0 ± 0.8 chiasmata per cell (Fig. 4). Notably, almost all analyzed *taf4b* lines exhibited a statistically significant reduction in crossover numbers, except for the *taf4b*_3_AA genotype (Fig. 4a; Kruskal-Wallis test, P < 0.0001, followed by Dunn’s post-hoc test). Within the *taf4b*_1 lineage, drops in chiasma counts were captured across genotypes, averaging 14.1 ± 0.8 for recessive homozygotes (aa), 12.8 ± 1.5 for heterozygous (Aa) individuals, and 14.0 ± 1 for dominant homozygotes (AA) (Fig. 4a).

**Figure 3.**
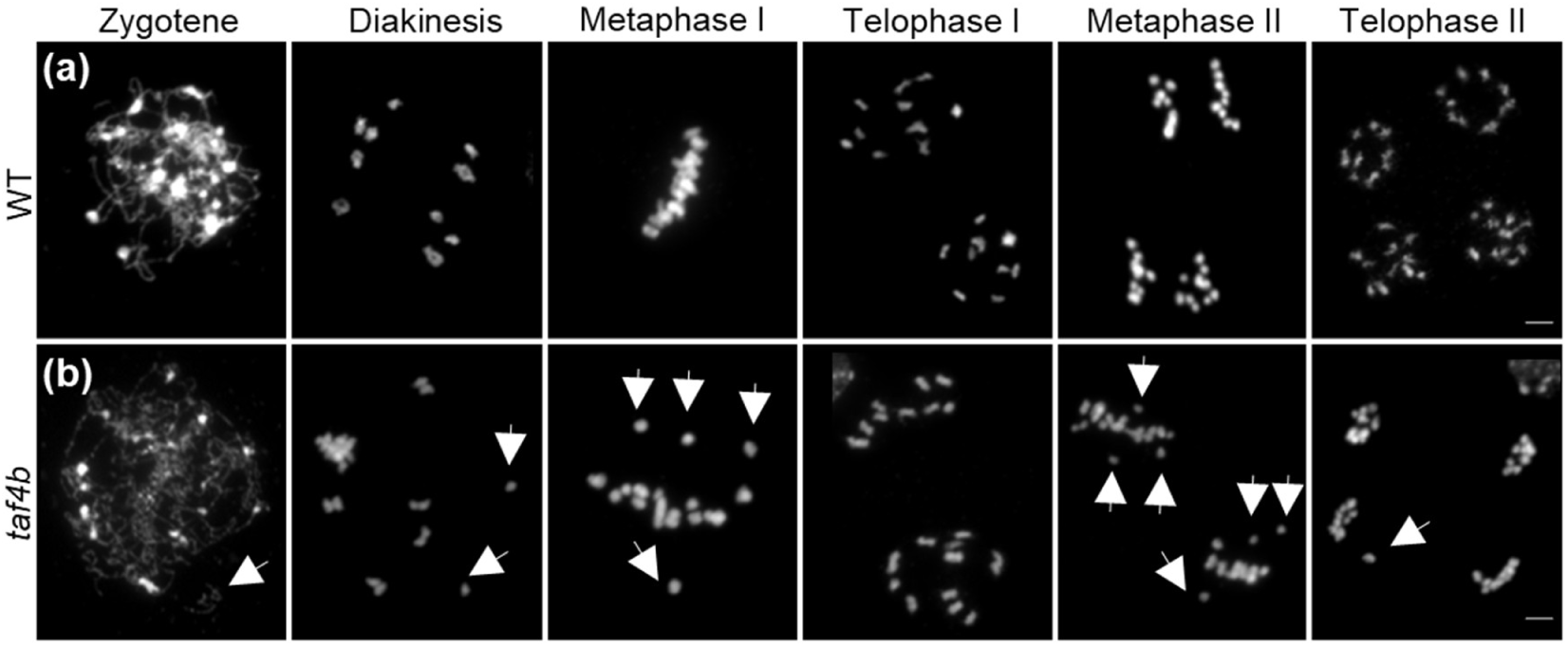
Male meiotic chromosome behavior in wild-type (WT) **(a)** and *taf4b* mutant plants **(b)**. White arrows indicate fragmented chromatin or univalents. DNA counterstained with DAPI is shown in gray. Scale bars: 5 µm

**Figure 4.**
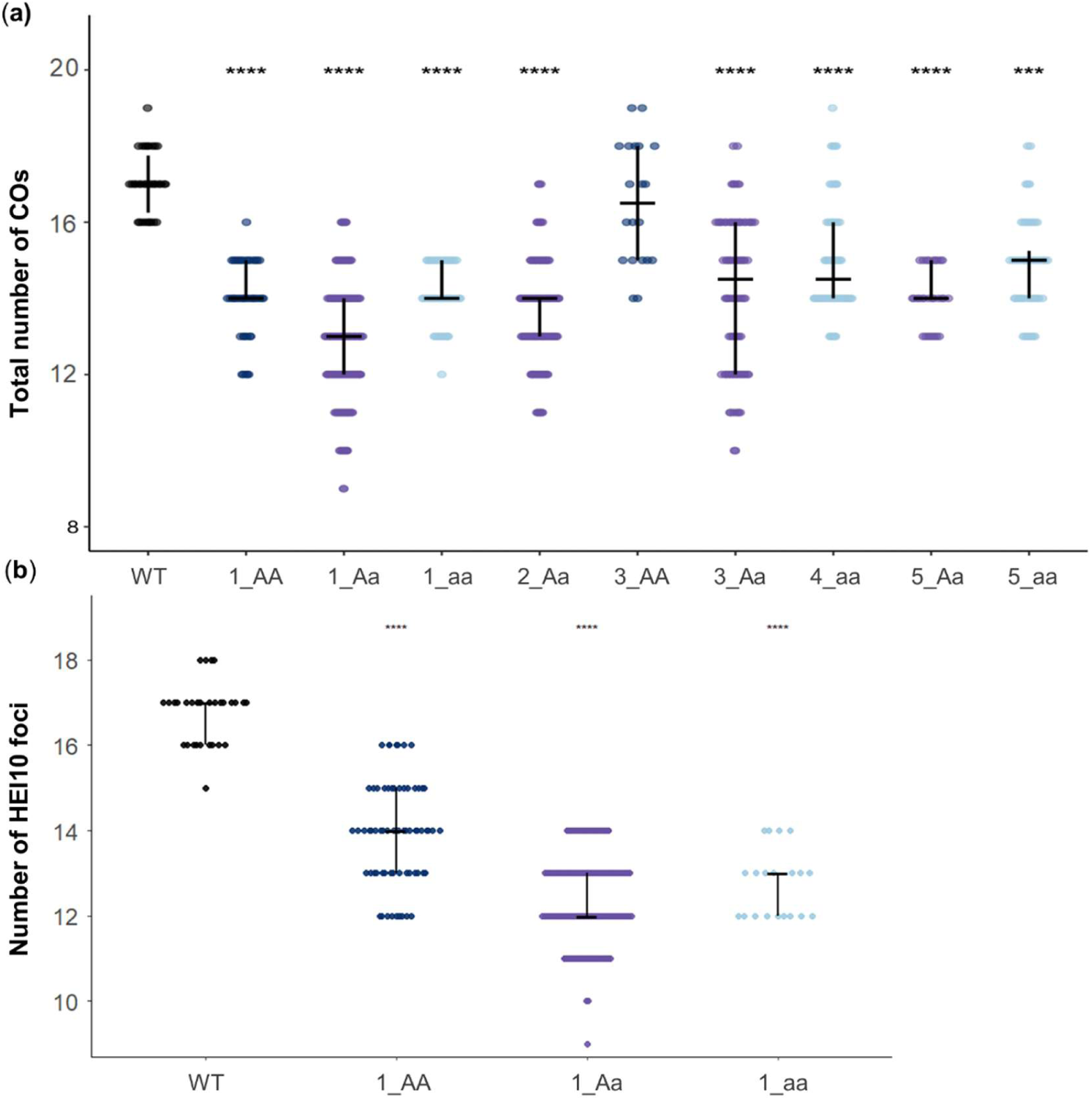
Evaluation of crossover frequency in *B. rapa* wild-type (WT) and *taf4b* mutant lines. (**a**) Crossover frequency estimated from chromosome configurations at diakinesis in WT and *taf4b* mutant lines in pollen mother cells (WT, (n = 34); *taf4b_*1_AA, (n = 45); *taf4b_*1_Aa, (n = 157); *taf4b_*1_aa, (n = 53); *taf4b_*2_Aa, (n = 124); *taf4b*_3_AA, (n = 20); *taf4b_*3_Aa, (n = 80); *taf4b_*4_aa, (n = 64); *taf4b_*5_Aa, (n = 29); and *taf4b_*5_aa, (n = 52) cells). Statistical significance was assessed using a Kruskal–Wallis test: ***, P < 0.001; ****, P < 0.0001. (**b**) Crossover frequency quantified by *HEI10* foci immunolocalization at the late pachytene stage in WT and the *taf4b* background (WT, (n = 30); *taf4b_*1_AA, (n = 66); *taf4b_*1_Aa, (n = 176); and *taf4b_*1_aa, (n = 20) cells).

Given the limited resolution of chiasma scoring, we independently assessed crossover designation using HEI10 immunolocalization. Since our comprehensive phenotypic screening demonstrated a robust and consistent reduction in chiasma frequency across the entire allelic series (Fig. 4a), the *taf4b*_1 line was selected as a representative genetic background (Fig. 4b; Fig. 5). While non-mutated wild-type control plants (R-o-18) displayed an average of 16.7 ± 0.7 HEI10 foci per cell, the *taf4b*_1 line as a whole showed a significantly reduced mean of 12.8 ± 1.2 foci per cell (Kruskal-Wallis test, P < 0.0001). To dissect the precise contribution of the *TAF4b* mutation relative to potential background variation, we analyzed individual segregating genotypes within this representative line (Fig. 4b). Recessive homozygotes (*taf4b*_aa) exhibited 12.8 ± 0.8 foci, heterozygotes (*taf4b*_Aa) showed 12.4 ± 1.0 foci, and dominant wild-type homozygotes (*taf4b*_AA) displayed 13.9 ± 1.2 foci.

**Figure 5.**
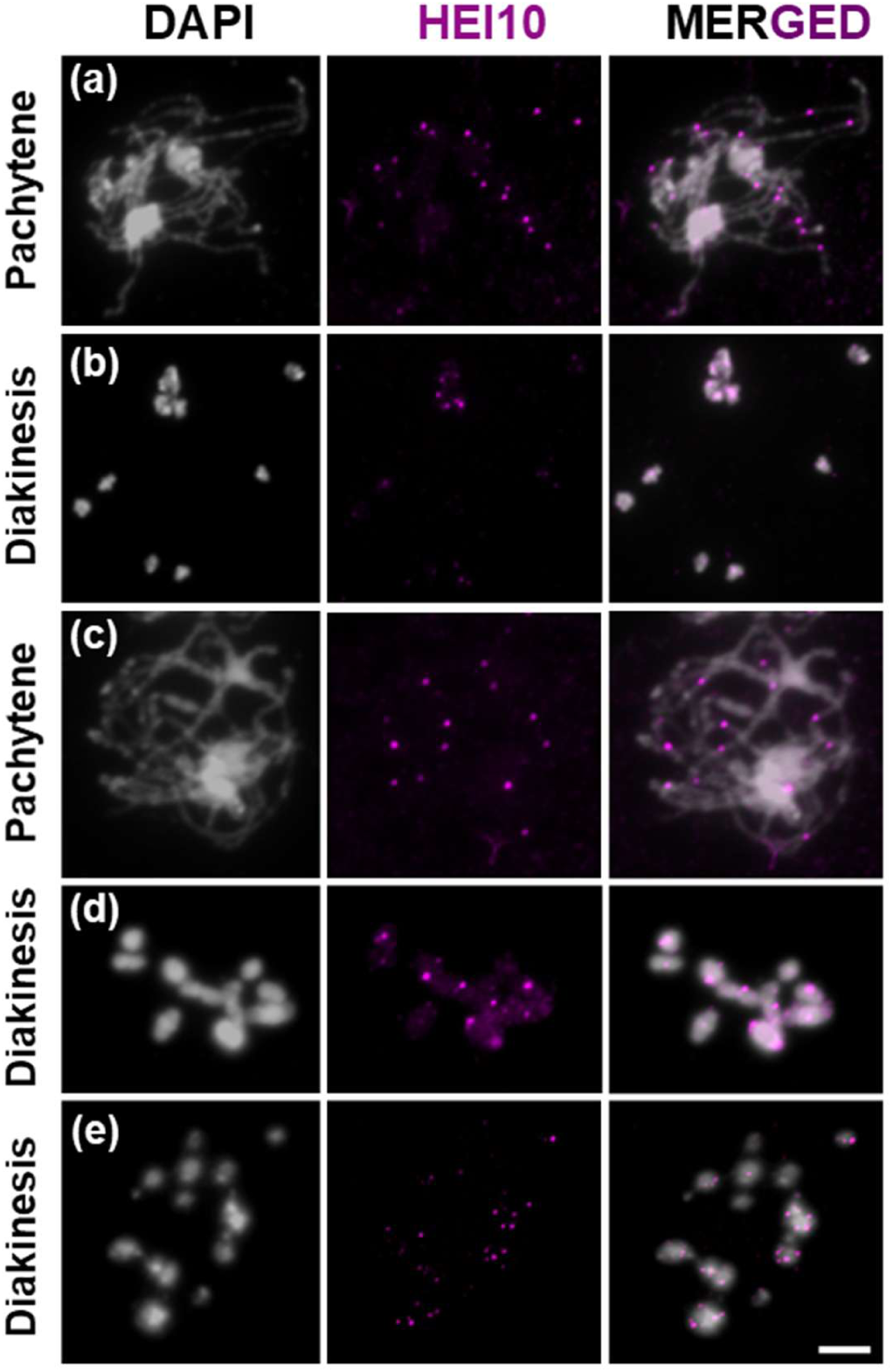
Class I crossovers (COs) visualized by HEI10 immunostaining in pollen mother cells. Panels show wild-type cells at pachytene **(a)**, and diakinesis **(b)**; and *taf4b*_1 mutant cells at pachytene **(c)** and diakinesis **(d, e)**. For the *taf4b*_1 mutant, two distinct diakinesis phenotypes are shown: a cell with ten intact bivalents **(d)** and a cell exhibiting a mix of bivalents and univalents **(e)**. Scale bar = 10 μm.

### Axis and synaptonemal complex loading were largely functional, with occasional chromatin detachment

To explore whether altered crossover frequency is associated with perturbations in prophase I chromosome structure, we performed immunostaining with ASY1 and ZYP1 antibodies to evaluate axis formation and synapsis in *taf4b*_1_Aa. Due to the limited availability of homozygous recessive mutant material, the cytological analysis was constrained to the heterozygous line. In wild-type meiocytes, synaptonemal complex (SC) formation progressed normally through prophase I: at leptotene, ASY1 localized along developing axial elements as thin filamentous signals; during zygotene, ZYP1 signals appeared between paired axes, consistent with synapsis initiation; and at pachytene, extended ZYP1 tracks marked synapsed homologs (Supplementary Fig. S2). In *taf4b*_1_Aa meiocytes, ASY1 and ZYP1 displayed largely normal axis establishment and ZYP1 loading. However, qualitative observations suggested a potential trend toward prolonged retention of the ASY1 antibody signal in a subset of mutant nuclei. Furthermore, distinct chromatin extensions, which appeared detached from the main chromatin mass, were occasionally observed during the zygotene–pachytene stages (Supplementary Fig. S2). This feature was not seen in wild-type, indicating occasional abnormalities in chromatin organization during prophase I in *taf4b*_1_Aa despite otherwise rather typical ASY1 and ZYP1 localization.

### Impairment of pollen viability and seed production in the *TAF4b* mutants

To test whether altered crossover frequency in *taf4b* lines correlated with compromised reproductive performance, we quantified pollen viability and seed set across the entire allelic series. Our phenotypic screening revealed a distinct structural-functional correlation depending on the mutation site relative to the predicted protein domains. While mutations residing within the variable N-terminal region (*taf4b*_2 and *taf4b*_3) maintained higher baseline fertility with noticeable individual variation, independent mutations disrupting the tightly conserved C-terminal linker cluster (*taf4b*_1, *taf4b*_4, and *taf4b*_5) consistently triggered a severe reduction in total reproductive success (Fig. 6).

**Figure 6.**
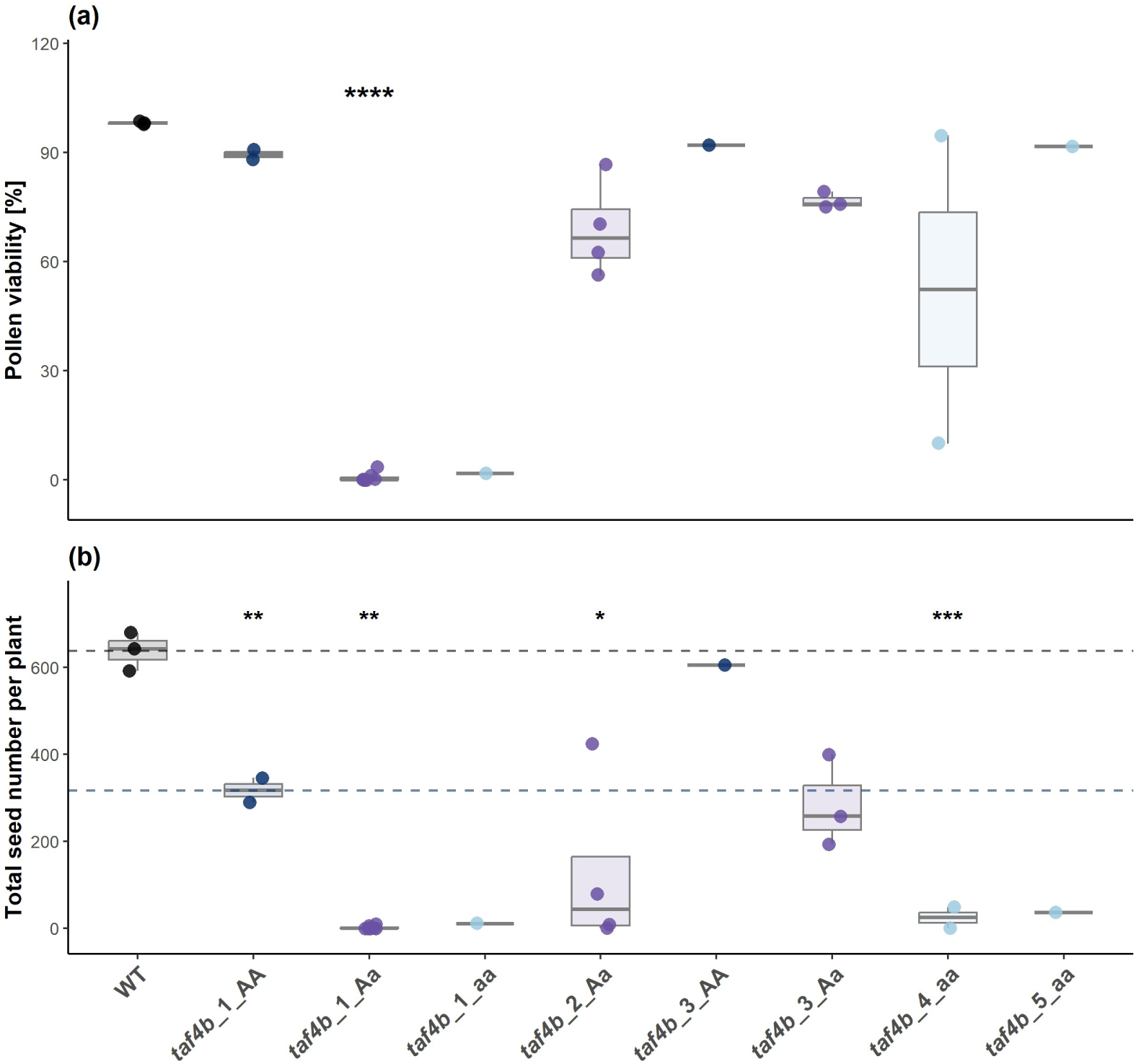
Assessment of reproductive fertility across *taf4b* mutant lines. **(a)** Pollen viability and **(b)** total seed number per plant collected after self-fertilization. Individual plants are represented as single dots. Dashed horizontal reference lines in panel **(b)** indicate the mean values for the wild-type (WT) control (black line) and the *taf4b_1_AA* background control (blue line). Statistical significance relative to the WT control was determined using a two-sided, two-sample Welch’s *t*-test (ns*, P* ≥ 0.05; *, *P* < 0.05; **, *P* < 0.01; ***, *P* < 0.001; ****, *P* < 0.0001).

Within the C-terminal cluster lineages, strong genotype-dependent effects were observed; for instance, heterozygous *taf4b*_1_Aa individuals showed near-complete male sterility, whereas dominant wild-type homozygotes (*taf4b*_1_AA) retained higher pollen viability (Fig. 6a). Crucially, while homozygous *taf4b*_4_aa and *taf4b*_5_aa plants retained high apparent pollen staining frequency, their total seed production dropped to near-zero levels, mirroring the severe seed-set failure of *taf4b*_1_aa (Fig. 6b). This marked discrepancy indicates that despite forming morphologically normal pollen grains, alleles targeting the C-terminal linker universally trigger a failure in gamete function or macrosporogenesis, suggesting an essential role for this conserved domain in *B. rapa* fertility.

To assess the genetic consistency of the TILLING lines and identify potential confounding background mutations, we performed whole-genome sequencing (DNA-seq) combined with transcriptomic profiling of background loci. After multiple rounds of stringent filtering, a total of 158 mutant-specific SNP sites were identified across the mutant lines, corresponding to 144 candidate genes with haplotype variation patterns matching that of the *taf4b_1* mutant plants (Supplementary Table S6). Among these background loci, two genes (*AXR1* and *FANCD2*) were associated with known meiosis-related functions, whereas several other loci, including *NOT1*, corresponded to genes implicated in pollen development. To evaluate whether these background mutations exhibited transcriptional activity that could interfere with the phenotypes, we examined the expression profiles of all 144 genes across our staged anther dataset. During meiotic interphase, only four background genes were differentially expressed between the mutant and the wild-type (two upregulated and two downregulated), and neither *AXR1* nor *FANCD2* exhibited significant differential expression (Supplementary Table S7). Similarly, none of the 144 background candidate genes showed significant transcriptomic alterations during the subsequent pachytene or telophase II stages. In somatic leaf tissues, four background genes were differentially expressed (one upregulated (AT3G08840) and three downregulated (*PICALM6*, *SD1-29*, and *ABCB11*), with only one gene overlapping with the interphase dataset in a reversed expression trend. These findings suggest that the expression of these background genes, including *AXR1* and *FANCD2*, remains largely unaltered, making them unlikely to confound the observed meiotic phenotypes.

### Transcriptomic profiling reveals stage-specific disruption at the onset of male meiosis

To determine whether the reduced crossover phenotype was caused by altered transcript abundance of the mutated gene itself, we first examined *TAF4b* expression levels across genotypes (WT, *taf4b*_1_AA, Aa and aa) and developmental stages. No significant differences in *TAF4b* expression were detected between any genotypes or time points, indicating that the phenotype is not driven by major changes in *TAF4b* transcription (Fig. 1). We next performed global differential expression analysis between the mutant lines (*taf4b*_aa and *taf4b*_Aa) and WT across staged anthers and leaf tissue, which revealed a pronounced stage dependence (Fig. 7). The most prominent transcriptional response (between *taf4b*_Aa/aa and _AA) occurred at interphase/leptotene, yielding 1 919 differentially expressed genes (DEGs; 1 138 downregulated and 781 upregulated, Supplementary Table S8), while 21 341 genes remained unchanged. In contrast, pachytene and telophase II showed minimal transcriptional changes (7 and 19 DEGs, respectively, Supplementary Table S9 and Table S10), whereas somatic leaf tissue displayed an intermediate response (702 DEGs; 184 downregulated and 518 upregulated, Supplementary Table S11).

**Figure 7.**
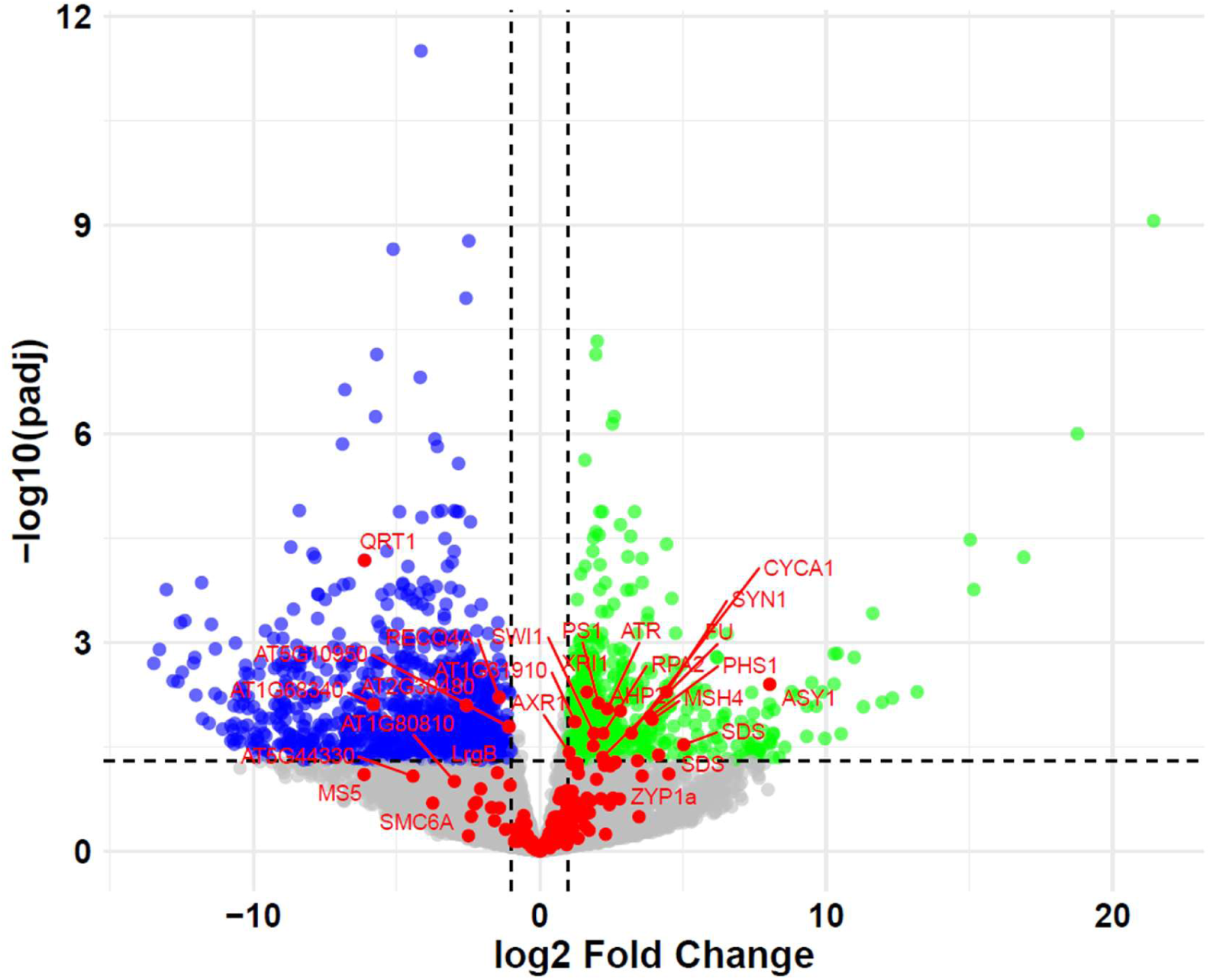
Volcano plot of differentially expressed genes (DEGs) between wild-type (WT) and the *taf4b* mutant. DEGs were identified at meiotic onset (interphase). The x-axis shows the absolute log₂ fold change (|log₂FC|), and the y-axis shows −log₁₀(adjusted P-value). Blue and green dots indicate genes significantly downregulated and upregulated in the *taf4b* mutant, respectively. Red dots highlight genes specifically associated with meiosis.

To further investigate this early disruption, we screened a curated set of 207 meiosis-related genes, revealing that 23 transcripts were significantly mis-regulated specifically at interphase/leptotene. These candidates exhibited a strong bias toward increased expression (19 upregulated, 4 downregulated, detailed information in Supplementary Table S8). The upregulated cohort included cohesion and axis/synapsis-associated factors (e.g., *REC8*, *ASY1*, *SWI1*, *PHS1*, *SDS*) as well as recombination, repair, and checkpoint components (e.g., *MSH4*, *RPA2A*, *ATR*, *HOP2/AHP2*). Conversely, downregulated loci included *RECQ4A*, the *HEI10*-interacting factor (*HEIP1*), and *PDS5*-like. This localized transcriptional dysregulation of essential chromosome pairing, recombination, and meiotic progression factors suggests a molecular mechanism contributing to the reduced crossover frequency observed in *taf4b*_1 mutant.

### Structural and biochemical features of the *taf4 a*llelic series: alternative splicing and TAF12 interaction dynamics

To investigate the potential structural and biochemical consequences of the identified amino acid substitutions within the C-terminal linker region, *in silico* post-translational modification profiling across the severe mutant alleles was performed (NetPhos 3.1). For the *taf4b*_1 (Ala515Thr) variant, the substitution was predicted to generate a novel, putative ectopic phosphorylation site as targeted specifically by Protein Kinase C (PKC). Similarly, profiling of the *taf4b*_5 (Pro499Ser) variant revealed the formation of two novel, distinct phosphorylation sites targeted by Casein Kinase I (CKI) and Protein Kinase A (PKA). In contrast, no novel phosphorylation sites were predicted for the *taf4b*_4 (Gly517Asp) substitution.

During sequence verification of cDNA clones generated for yeast two-hybrid assays, we identified multiple *TAF4b* transcript variants consistent with alternative splicing (six obtained by sequencing 18 clones). These variants were recovered from leaf-derived clones exclusively in the recessive mutant (*taf4b*_1_aa) background (Supplementary Fig. S3). The detected differences predominantly affected exon 2 and exon 5, indicating that these regions contribute to transcript diversity. In all cases, the open reading frame was maintained, consistent with the potential production of distinct protein isoforms. Importantly, although alternative transcripts were not explicitly assembled from our reproductive datasets, the inherent resolution limits of short-read RNA-seq precluded the definitive exclusion of these *TAF4b* splice variants in meiotic tissues.

To evaluate whether the core molecular function of TAF4b is affected by the mutation, we performed pairwise yeast two-hybrid (Y2H) assays. The cDNA clone selected for the Y2H assays was specifically chosen to reflect the precise splice structure of the *TAF4b* transcript identified in our reproductive anther sequencing data, ensuring its physiological relevance to the meiotic program. Prior to interaction testing, expression of the three bait proteins (TAF12, TAF4b-WT, and TAF4b-M) was confirmed to be non-toxic, as yeast transformants exhibited growth rates comparable to the empty pDEST32 vector control on SC-Leu medium (Fig. 8a). Furthermore, negative controls demonstrated only marginal autoactivation across all reporter selection media, including SC-Leu-Trp-Ura, SC-Leu-Trp-His + 3-AT, SC-Leu-Trp + 5-FOA, and X-gal plates (Fig. 8b).

**Figure 8.**
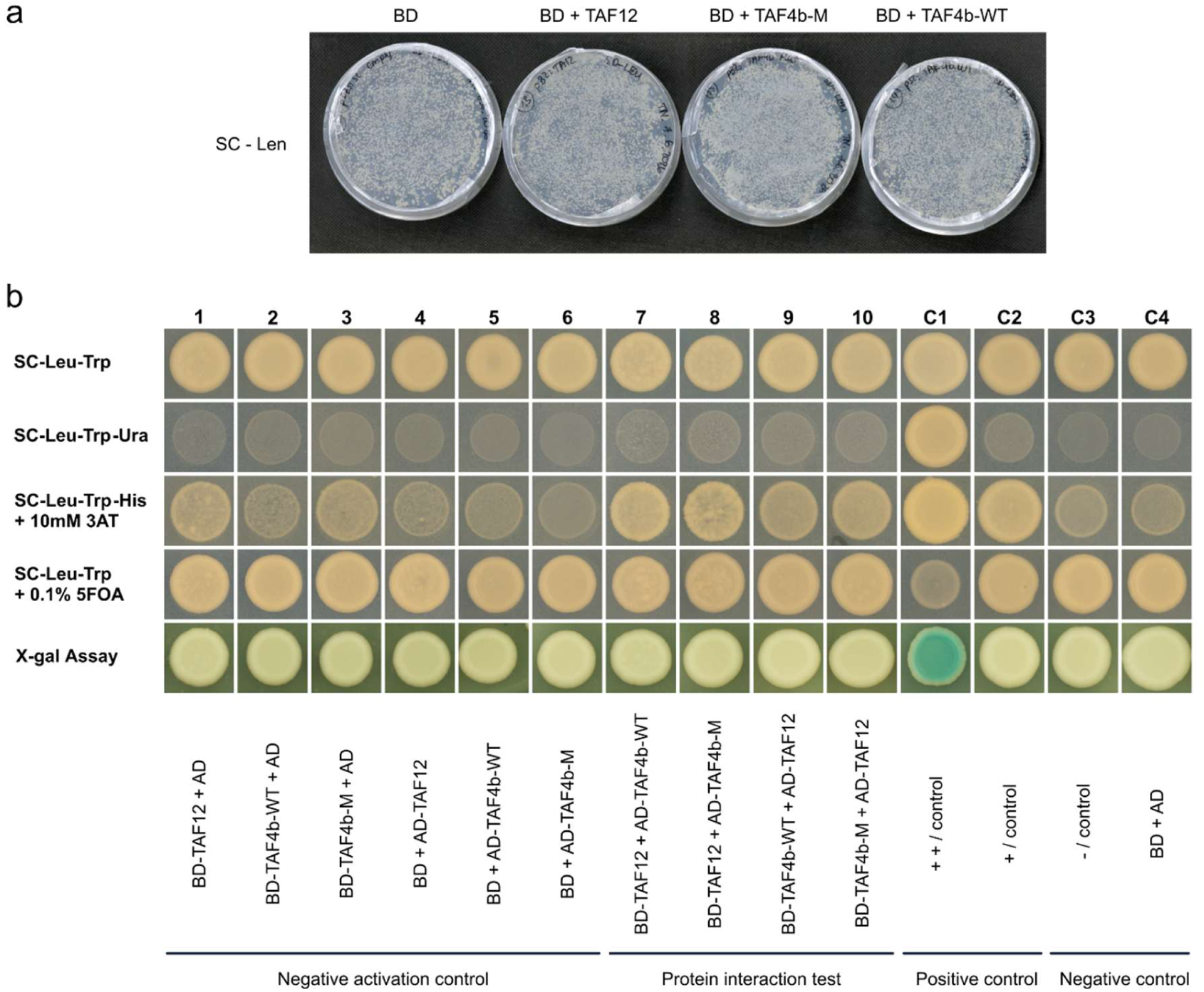
Pairwise yeast two-hybrid (Y2H) analysis of TAF4b and TAF12 interactions. (a) Toxicity testing of TAF12, TAF4b-M, and TAF4b-WT bait proteins. Transformants exhibit comparable growth rates to the empty pDEST32 vector control, indicating a lack of protein toxicity; (b) Pairwise Y2H interaction assays. Columns 1–6 show autoactivation assays for the respective bait and prey constructs. Columns 7–10 display reciprocal interaction tests between TAF4b (WT or M) and TAF12. Controls include: C1, strong positive control; C2, weak positive control; C3, negative interaction control; and C4, negative autoactivation control (empty pDEST32 and pDEST22 plasmids).

Under stringent selection setups (SC-Leu-Trp-Ura, SC-Leu-Trp + 0.1% 5-FOA, and X-gal), reporter gene activation was restricted exclusively to the strong positive control, with no detectable signals in the test samples. However, under moderated selective conditions utilizing SC-Leu-Trp-His supplemented with 10 mM 3-AT, yeast cells expressing the test combinations revealed that wild-type *B. rapa* TAF4b (TAF4b-WT) establishes a weak but reproducible physical interaction with TAF12, consistent with its expected structural role within TAF-containing transcription initiation complexes. Notably, the mutant TAF4b-M protein retained a comparable level of interaction under these specific selective conditions. This indicates that the identified single-nucleotide polymorphism does not operate via a simple disruption or total loss of TAF12 subunit binding (Fig. 8b).

## Discussion

In this study, we establish the unique *TAF4b* locus on chromosome A7 as a non-redundant transcriptional gatekeeper required to program early prophase I entry and ensure meiotic crossover assurance in *Brassica rapa*. We found that a significant reduction in chiasma frequency was a universal feature across the analyzed *taf4b* genotypes. Notably, subsequent reproductive fertility defects exhibit a clear structural-functional partition: while N-terminal variants (*taf4b*_2 and *taf4b*_3) maintain higher reproductive stability, mutations targeting the highly conserved C-terminal linker region (*taf4b*_1, *taf4b*_4, and *taf4b*_5) drive a consistent and severe reduction in total seed set.

Through the integration of genomic background sequencing, quantitative cytogenetics, and transcriptomic profiling, we show that this core meiotic disruption is driven by *taf4b* dysfunction and associated with a stage-specific transcriptional dysregulation at the onset of prophase I. Together with interaction evidence indicating that the *taf4b*_1 mutation impairs meiotic progression without completely abolishing TAF12 subunit binding, these findings provide comprehensive insights into how a core transcription factor coordinates stage-specific gene expression to ensure efficient meiotic recombination.

The universal reduction in chiasma frequency and HEI10 foci we observed suggests *TAF4b* is an evolutionarily-conserved factor for robust crossover formation in *B. rapa*. However, the severity of this reduction varied between lines, suggesting that the magnitude of the meiotic defect is related to the functional severity of the modified variant relative to its structural domain positioning, where alleles striking the conserved C-terminal linker region induce a significantly more severe reproductive impairment than those disrupting the variable N-terminal segment. Such variations, where the structural position or abundance of specific transcripts dictates the severity of phenotypic outcomes, have been well-documented in other plant meiotic genes. For example, *Arabidopsis thaliana SPO11-1* transcript dosage directly calibrates DSB levels (Xue et al. 2018), while alterations in functional *MSH4* copy number and expression in *Brassica napus* trigger extensive univalent formation (Gonzalo et al. 2019).

The emergence of univalents and increased rod bivalents indicates a qualitative crossover assurance failure across severe alleles, leading to achiasmate chromosome segregation (Fig. 3; Fig. 4a). Unlike *A. thaliana*, where *taf4b* mutants where *taf4b* mutants completely lack univalents and preserve wild-type silique seed counts (Lawrence et al. 2019), *B. rapa* demonstrates a distinct evolutionary requirement for this single-copy subunit. This fertility dependence parallels the essential role of *TAF4b* in mammals, where its depletion causes germ cell loss and sterility (Freiman et al. 2001; Falender et al. 2005; Grive et al. 2016). Although chemical mutagenesis inherently introduces confounding background mutations, the extensive genomic background filtering employed during our line establishment, combined with the precise transcriptional dysregulation of core meiotic machinery observed via RNA-seq, strongly suggests that this phenotype is a direct consequence of the *taf4b* lesions rather than off-target genetic effects. This severe phenotype may reflect species-specific thresholds in recombination buffering capacity, where *TAF4b* dysfunction drops recombination-related functions below a critical limit, positioning *TAF4b* upstream as an operator of the early prophase I program.

The evolutionary divergence between *TAF4* and *TAF4b* genes in the *B. rapa* genome highlights a potential functional specialization shaped that predates the *Brassica* lineage, which was subsequently maintained during post-polyploidization genome remodeling (Fig. 6). While the duplicated *TAF4* loci on chromosomes A2 and A6 likely support somatic functions, our expression profiling revealed that the unique *TAF4b* gene on chromosome A7 exhibits a distinct, meiosis-enriched expression pattern (Fig. 1). This highly specific recruitment explains why mutations in the A7 locus trigger a severe meiotic phenotype without functional compensation by the related somatic *TAF4* copies. Biochemical and structural eukaryotic data have established that TAF4/TAF4b subunits directly heterodimerize with TAF12 via conserved histone-fold domains to maintain core TFIID transcription complex integrity (Werten et al. 2002; Gazit et al. 2009). To evaluate how the specific *taf4b_1* missense mutation impacts this essential molecular association, we performed yeast two-hybrid (Y2H) assays, which showed that both the wild-type TAF4b and the mutant protein interact weakly with TAF12 (Fig. 8). Within TFIID, such low-affinity dynamics are common and often depend on additional scaffolding factors *in vivo* (Wright et al. 2006). Crucially, because the *taf4b_1* mutation did not disrupt this baseline interaction, the severe meiotic defects cannot be attributed to a simple loss of TAF12 assembly. Instead, these phenotypic disruptions likely stem from subtler downstream regulatory alterations affecting core promoter engagement.

*In silico* structural profiling provides biochemical insights to interpret the severe reproductive and meiotic phenotypes unique to the C-terminal linker cluster mutations. The Ala515Thr substitution in *taf4b_1* and the Pro499Ser substitution in *taf4b_5* are predicted to generate novel, ectopic phosphorylation sites, recognized by standard cellular kinases, including PKC, CKI, and PKA. Concurrently, while the Gly517Asp substitution in *taf4b_4* does not introduce novel phosphorylation targets, the replacement of a neutral glycine with a negatively charged aspartic acid residue represents a classic phosphomimetic mutation. Together, these data strongly suggest that introducing a negative charge cascade within this highly conserved linker region, either via active ectopic phosphorylation or structural mimicry, alters TAF4b stability, induces destructive conformational changes, or compromises the recruitment of meiotic partners during early prophase I. This model aligns with evidence that the TAF4/TAF4b–TAF12 module contributes to core promoter engagement, where perturbing structural conformation diminishes TFIID occupancy at specific promoters (Gazit et al. 2009). Because plant gene promoters often coincide with meiotic DSB and crossover hotspots (Choi et al. 2018; Serra et al. 2018), such localized electrostatic alterations within the TFIID complex could directly disrupt chromatin accessibility for core recombination machinery across the entire C-terminal cluster.

The identification of multiple *TAF4b* splice variants in our material points to an additional layer of transcript complexity in *B. rapa* (Supplementary Fig. S3). Alternative splicing of human *TAF4* functionalizes distinct isoforms (Kazantseva et al. 2013, 2016), suggesting a similar transcriptional processing mechanism may apply in the plant context. In our study, recurrent alterations in exon 2 and exon 5 preserved the open reading frame, consistent with the production of structurally distinct protein isoforms. While these variants were initially identified via cloning in somatic leaf tissue, our transcriptomic sequencing and genomic visualization across both vegetative and reproductive structures confirmed their presence in both leaves and isolated meiocytes. Notably, no statistically significant differences in isoform expression levels were detected between the wild-type and *taf4b*_1_aa backgrounds. This baseline transcript versatility across distinct cell lineages indicates that the alternative splicing of *TAF4b* represents a constitutive, tissue-independent feature of its expression rather than a mutant-specific disruption or a somatic-only phenomenon. Consequently, these findings highlight the inherent complexity of TAF4b post-transcriptional regulation during both vegetative development and generative phases.

Our cytological findings indicate that the major structural steps of prophase I chromosome remodelling - axis establishment and SC loading - proceed largely normally in *taf4b*_1_Aa male meiocytes. Immunostaining of ASY1 along chromosomes and extended ZYP1 tracks during zygotene–pachytene were broadly comparable to the wild-type, arguing against a pervasive failure in axis formation or bulk SC polymerization (Supplementary Fig. S2). While this structural integrity matches our RNA-seq data showing that core axis/SC transcripts are not depleted, normal plant meiosis requires progressive depletion of ASY1 from chromosome axes as synapsis proceeds (Armstrong et al. 2002; Yang et al. 2022). Interestingly, our qualitative cytological observations revealed a trend toward prolonged ASY1 retention, suggesting a delay in axis disassembly. This phenotype aligns with the transcriptomic upregulation of ASY1 in our early meiotic dataset, a phenomenon known to disrupt SC dynamics and axis remodeling when core component stoichiometry is altered (Lambing et al. 2020).

Against this background of functional loading, a subset of zygotene–pachytene nuclei exhibited a mutant-specific phenotype of thread-like chromatin extensions detached from the main chromatin mass. These localized aberrations coincide with the window when recombination intermediates are processed and crossover sites designated. In plants, the SC actively modulates crossover regulation, where perturbations in chromosome/SC context deeply influence crossover assurance and spacing (France et al. 2021). Recent work emphasizes that HEI10*-*dependent patterning reflects dynamic processes along synapsed chromosomes sensitive to underlying structure (Morgan et al. 2021). Accordingly, our immunolocalization data showed that HEI10 foci are significantly reduced in *taf4b* meiocytes. Because HEI10 chromosome-associated foci provide a reliable readout for class I crossover designation (Chelysheva et al. 2012), this reduction demonstrates a depletion of crossover sites. Together, these observations support a model in which *TAF4b* is not required for gross axis or SC loading, but is essential for robust recombination pathway progression. This cytological depletion of HEI10 likely reflects the early transcriptional dysregulation captured via RNA-seq, suggesting that *TAF4b*-mediated gene expression at meiotic onset is required to establish downstream crossover designation and prevent the univalents observed in *taf4b*_1.

Our global RNA-seq profiling revealed that the *taf4b* dysfunction causes a pronounced transcriptional shift specifically at interphase/leptotene (∼1919 DEGs), whereas later meiotic stages show relatively few changes (Fig. 7). This striking stage specificity points to a pivotal role for *TAF4b* in establishing or buffering the transcriptional state required for meiotic entry. At the interphase/leptotene transition, we observed altered expression of core axis, synapsis, and cohesion regulators (e.g., *REC8*, *ASY1*, *SWI1*, and *PHS1*), alongside components linked to early recombination processing and genome surveillance (e.g., *RPA2A*, *BRCA2B*, *ATR*, and *PS1*). These patterns suggest that an early transcriptomic imbalance can directly perturb chromosome architecture and the cellular response to recombination intermediates, modifying downstream crossover outcomes. Comparison with *A. thaliana* supports the idea that *TAF4b* broadly shapes meiotic gene expression, where its loss triggers widespread transcriptomic changes in purified meiocytes, confirming that *TAF4b* supports a dedicated germline transcriptional program (Lawrence et al. 2019). Similar “early-entry” sensitivities and tight transcriptional checkpoints are well-documented across higher plants. In maize (*Zea mays*), *AMEIOTIC1* (*AM1*) is strictly required for meiotic entry, illustrating that early prophase I progression depends on dedicated regulators coordinating sequential chronological events (Pawlowski et al. 2009). In *A. thaliana*, early prophase I is likewise highly dependent on cohesion and axis regulation: for instance, *SWI1/DYAD* antagonizes *WAPL* to maintain meiotic cohesin, directly linking cohesion status to recombination outcomes (Yang et al. 2019). Furthermore, transcriptional control of meiotic progression has clear precedents, such as the chromatin-remodeling protein *DUET/MMD1*, which directly regulates meiotic expression and cell-cycle transitions (Andreuzza et al. 2015). Together, these data indicate that the *taf4b* mutation primarily perturbs transcriptional programs at the onset of male meiosis, prior to or coincident with early prophase I events, thereby placing *TAF4b* upstream of the core recombination machinery. Disruption of this early program provides a plausible molecular route to the reduced numbers of ring bivalents and occasional univalents observed, because early perturbations in axis/cohesion formation and checkpoint balance can shift DSB repair away from class I crossover designation.

Beyond meiotic recombination, our *taf4b* lines exhibited reduced pollen viability and seed set, showing a profound structural dependence based on the position of the mutation. Transcriptomic downregulation of numerous tapetum and sporopollenin/exine pathway genes during interphase/leptotene suggests that defects in anther somatic support programs contribute to this male sterility (Ariizumi & Toriyama, 2011; Shi et al. 2015). This phenotype contrasts with *A. thaliana*, where *taf4b* mutants maintain wild-type seed numbers per silique despite crossover reductions (Lawrence et al. 2019). Although background mutations inherent to TILLING resources, such as the missense lesion identified here in the pollen-development factor *NOT1* (Lin et al. 2020), likely contribute to the specific male gametophyte abnormalities observed in *taf4b_1* (Fig. 6a), they do not account for the global reproductive failure. The fact that independent C-terminal alleles (*taf4b*_4_aa and *taf4b*_5_aa) uniformly manifest an identical severe reduction in seed set, combined with the wild-type fertility retained by the *taf4b*_3_AA background control (∼605 seeds), demonstrates that the core sterility phenotype is driven by *taf4b* dysfunction.

The unique reproductive discrepancy captured in *taf4b*_4_aa and *taf4b*_5_aa, where high pollen staining frequency coexists with near-complete sterility, indicates that these specific missense mutations preserve initial pollen wall development but compromise functional gamete quality or maternal macrosporogenesis. These early imbalances in core structural components likely yield aneuploid gametes that ultimately fail during post-meiotic developmental checkpoints. Consequently, this discrepancy indicates that despite producing morphologically normal pollen, alleles targeting the C-terminal linker disrupt gamete function or macrosporogenesis. This highlights a novel post-polyploidization specialization for *B. rapa TAF4b* as a critical gatekeeper of both meiotic crossover assurance and reproductive success.

## Acknowledgements

We would like to thank Mathilde Grelon (Université Paris-Saclay, INRAE) and Raphael Mercier (Max Planck Institute for Plant Breeding Research) for generously sharing the HEI10 antibody, and Li Guo (University of Bonn) for valuable advice regarding the yeast two-hybrid assays.

## Author contributions

JM designed the research, performed the experiments, analyzed the data, and wrote the manuscript. RD managed the plant material and assisted with media preparation for Y2H assays. ZL analyzed and visualized the DNAseq and RNAseq data, and contributed to manuscript writing. TDN designed and performed molecular cloning and Y2H experiments, analyzed the alternative splicing data, prepared the figures for both Y2H and alternative splicing analyses and contributed to manuscript writing. JS performed molecular cloning and Y2H experiments. ASM designed the research, acquired funding, and contributed to the critical revision of the manuscript. All authors read and approved the final manuscript.

## Data availability

DNA/RNA sequencing data were deposited at European Nucleotide Archive (ENA; https://www.ebi.ac.uk/ena/browser/home) PRJEB125661. All other relevant data can be found within the manuscript and its supporting materials.

## Supplementary information

**Supplementary Table S1.** Transcriptome mapping statistics and unique mapping ratios.

**Supplementary Table S2.** Whole genome sequence mapping statistics and unique mapping ratios.

**Supplementary Table S3.** Primers used for molecular cloning in this study. F, forward primer; R, reverse primer.

**Supplementary Table S4.** Combinations of bait and prey plasmids co-transformed into yeast strain *MaV203*.

**Supplementary Table S5.** Expression levels and statistical comparisons of *TAF4b* (A7) and *TAF4* (A2 and A4) across materials and developmental stages.

**Supplementary Table S6.** Variant sites and functional annotations of the 144 candidate genes identified by whole-genome resequencing.

**Supplementary Table S7.** Expression profiles of DEGs among the 144 candidate genes identified by whole-genome resequencing.

**Supplementary Table S8.** All expression profiles and functional annotations between mutant (*taf4b*_aa and *taf4b*_Aa) vs. WT (*taf4b*_AA) during interphase.

**Supplementary Table S9.** All expression profiles and functional annotations between mutant (*taf4b*_aa and *taf4b*_Aa) vs. WT (*taf4b*_AA) during pachytene.

**Supplementary Table S10.** All expression profiles and functional annotations between mutant (*taf4b*_aa and *taf4b*_Aa) vs. WT (*taf4b*_AA) during telophase.

**Supplementary Table S11.** All expression profiles and functional annotations between mutant (*taf4b*_aa and *taf4b*_Aa) vs. WT (t*af4b*_AA) during leaf.

---

**Supplementary Figure S1.**
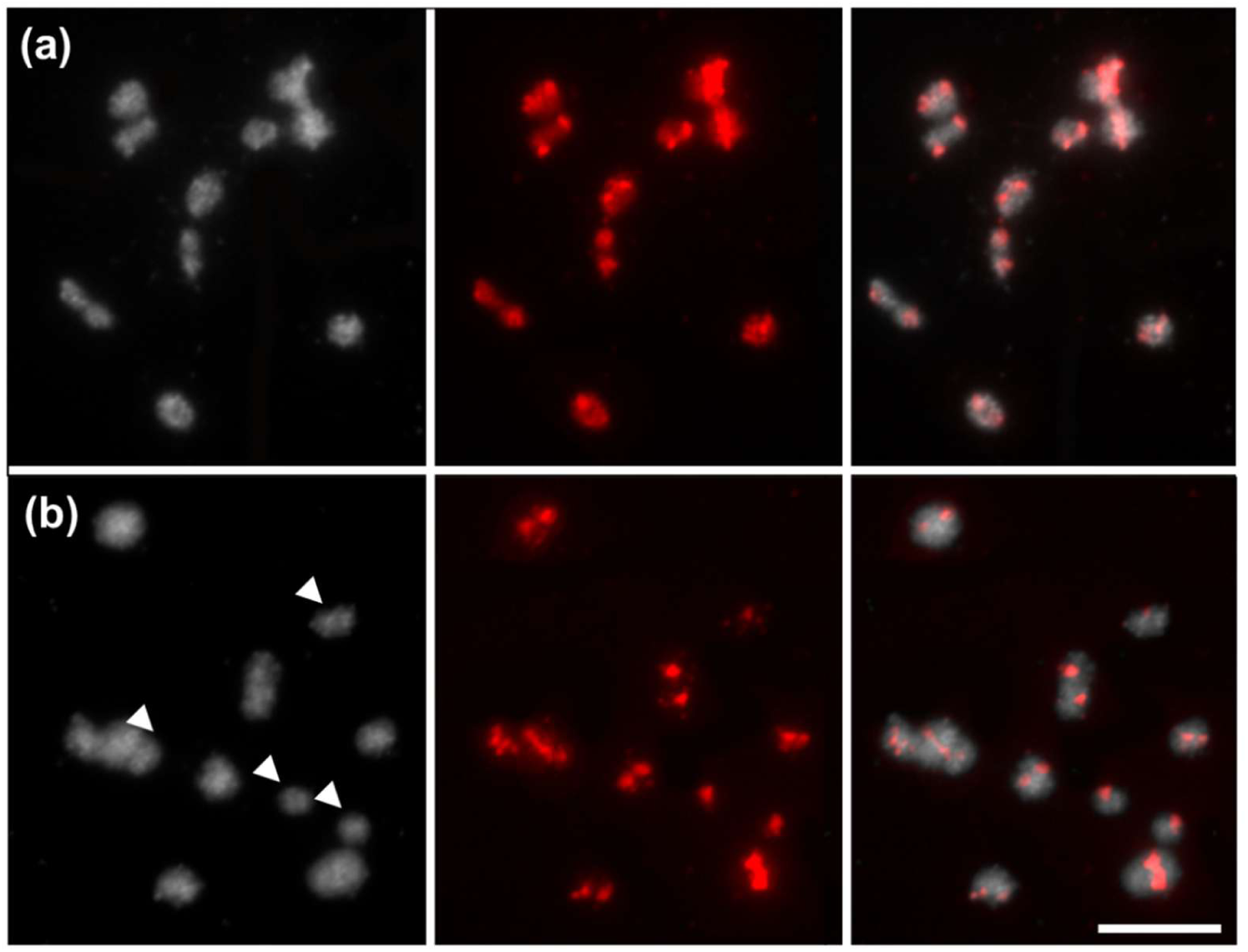
Validation of univalent formations via centromere-specific fluorescence *in situ* hybridization (FISH). (a) Pollen mother cell from a wild-type (WT) control plant at diakinesis showing ten intact, regular bivalents. (b) Pollen mother cell from the *taf4b_1* mutant line displaying a mixture of bivalents and univalents (arrowheads). Chromosomes were counterstained with DAPI (gray); centromeric probe (red). Scale bar = 5 µm.

**Supplementary Figure 2.**
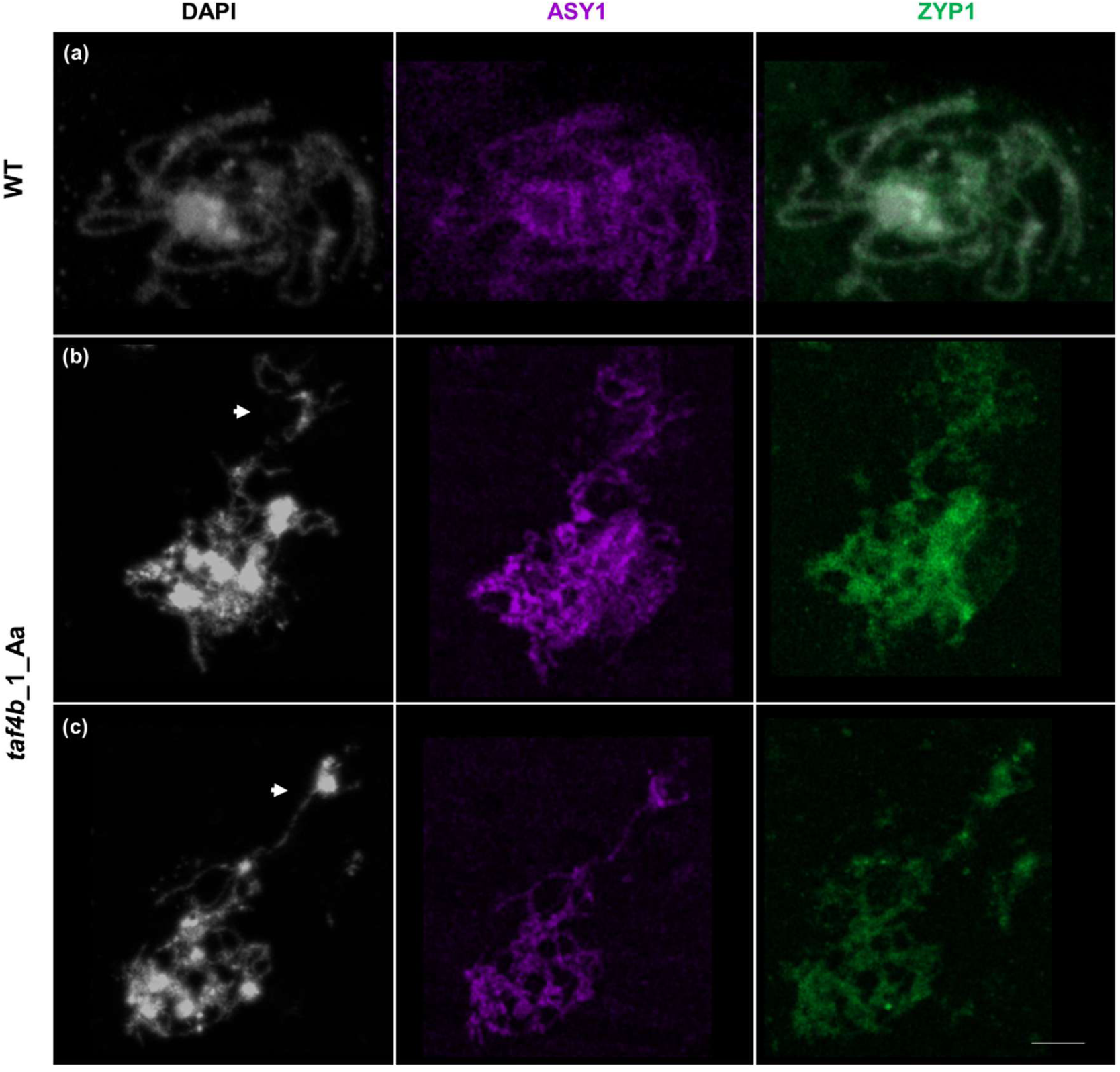
Immunostaining with ASY1 and ZYP1 antibodies at pachytene in male meiosis of WT **(a)** and *taf4b*_1_Aa mutant plant **(b, c)**. White arrows indicate chromatin threads detached from the main chromatin mass. ASY1 (purple); ZYP1 (green), chromatin (gray). Scale bar 5 μm.

**Supplementary Figure 3.**
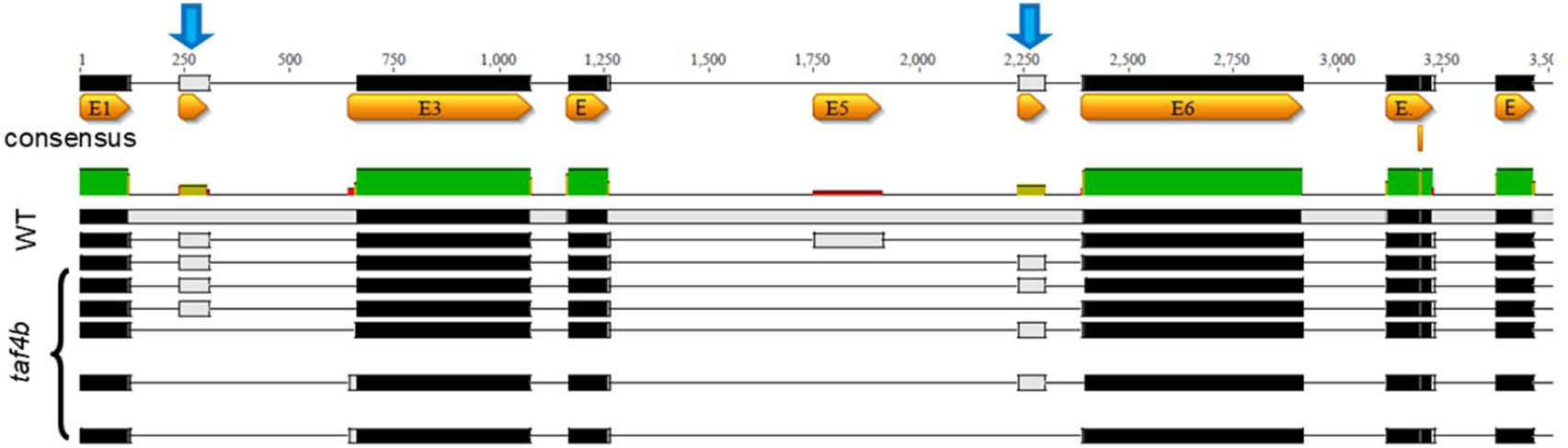
Alternative splicing variants of *TAF4b*. Schematic representation of the *TAF4b* gene and transcript structure in *Brassica rapa* wild-type (WT, R-o-18) and the recessive *taf4b*_1_aa mutant. Boxes and lines represent exons and introns, respectively. *TAF4b*-gDNA and *TAF4b*-cDNA serve as reference sequences. The WT background exhibits a single predominant transcript isoform (*TAF4b*_WT), whereas six distinct alternative splicing variants were identified from 18 sequenced cDNA clones in the *taf4b*_1 mutant line. Diagram generated using Geneious Prime.

